# Patch-leaving decisions and pupil-linked arousal systems

**DOI:** 10.1101/2024.09.06.611272

**Authors:** Anna Marzecová, Brent Vernaillen, Drita Hoxhaj, Luca F. Kaiser, Tom Verguts, Gerhard Jocham

## Abstract

Deciding when to abandon a depleting resource in favour of potentially richer alternatives is fundamental to adaptive behaviour. Such patch-leaving decisions require balancing the expected advantage of leaving against both the cost of moving and the reward foregone in the current environment. Previous research suggests that activity of noradrenergic (NE) neurons in the locus coeruleus (LC) underpins patch-leaving. In the current study, we used pupil dilation as a time-resolved readout of subcortical neuromodulation during patch-leaving. We hypothesised that leave decisions will be preceded by a transient pupil dilation. Participants harvested from exponentially depleting patches (blueberry bushes) with three initial reward values, in two environments, which differed in the variability of initial reward values. Behavioural results show that, as predicted by the mathematically optimal solution, participants adjusted their decisions based on the instantaneous reward rate, but also displayed a bias to overharvest (stay longer in compared to the optimum), which was more pronounced in the high variability environment. Pupil size was overall larger in the high variability environment, associated with increased uncertainty. Importantly, we observed an increased transient pupil dilation in response to reward outcomes immediately preceding leave compared to stay decisions, as well as an increase in pupil dilation (and RT) across successive stay trials, leading up to leave decisions, presumably indicating an increased LC-NE activity associated with abandoning current options to explore alternatives.

## Introduction

Deciding when to abandon a depleting resource in favour of potentially richer alternatives is fundamental to adaptive behaviour. Rewards in the natural environment are not distributed uniformly, but are contained in areas (patches), which deplete over time when harvested (Stephens & Krebs, 1986). Successful foraging for resources requires balancing the expected advantage of leaving the currently harvested patch for another one against both the cost of travelling to another patch, and the reward foregone in the current patch. These serial stay-or-leave dilemmas, coined as “patch-leaving” decisions, are ubiquitous in daily life, from food search to decisions on whether to set out on the effortful move to another country in pursuit of more rewarding opportunities. Notably, the serial dependency inherent to patch-leaving decisions sets them apart from value-based choices, thought to be reflected in different neural correlates of value-based vs patch-leaving choices (Constantino et al., 2017; Kaiser et al., 2021; Kolling et al., 2012).

The mathematically optimal solution to the patch-leaving problem is given by the marginal value theorem (MVT; Charnov, 1976), which stipulates that an agent should leave the patch when the instantaneous reward rate falls below the average reward rate in the environment. As a result, the agents leave earlier from patches with lower instantaneous reward rate compared to patches with higher reward rate. While the MVT successfully describes behaviour across species (Charnov, 1976; Constantino & Daw, 2015; Hayden et al., 2011; Kane et al., 2019), it has also been shown that individuals commonly stick to harvesting from a current resource longer than optimal, a phenomenon termed overharvesting or overstaying; (Kane et al., 2019; Kendall & Wikenheiser, 2022).

The computations underlying patch-leaving decisions have been proposed to rely on the activation of several neuromodulatory pathways (Calhoun & Hayden, 2015; Marzecová et al., 2021), with a focus on catecholaminergic systems (Bendesky et al., 2011; Hills, 2006). In particular, the locus coeruleus-noradrenergic (LC-NE) system, with its key role in regulating adaptive and flexible behaviour (Poe et al., 2020), is thought to be also involved in the control of patch-leaving behaviour (Cremer et al., 2022; Doren et al., 2023; Kane et al., 2017). The adaptive gain theory (AGT, Aston-Jones & Cohen, 2005) describes two modes of LC activity associated with distinct arousal and behavioural states: a tonic firing mode with sustained activation linked to disengagement from the task (exploration), and a phasic mode with high frequency bursts in response to task-relevant stimuli, facilitating task-engagement and action execution (exploitation). Another theoretical approach proposes that transient bursts of LC activity following outcomes lead to reorganisation of functional networks (‘network reset’, (Bouret & Sara, 2005; Sara & Bouret, 2012) allowing shifts in cognitive representations and behavioural adaptations (McBurney-Lin et al., 2022; Ogg et al., 2023).

The empirical evidence on the involvement of LC-NE in patch-leaving is scarce. In rodents, chemogenetic stimulation of LC tonic activity has been shown to impair patch-leaving, with early patch-leaving due to increased decision noise (Kane et al., 2017). In humans, a recent study inhibiting NE reuptake using reboxetine suggests lower patch-leaving thresholds with higher NE levels (Doren et al., 2023). Given the distinct roles of phasic and tonic modes and their inverse relationship (Li et al., 2023), the dynamics of the transient LC-NE indexed by trial-by-trial fluctuations during patch-leaving decisions is yet to be described.

In the current study, we utilised pupil dilation as a non-invasive, time-resolved readout of subcortical neuromodulation during patch-leaving. Pupil size has been shown to be partially under the control of locus coeruleus NE neurons (Joshi & Gold, 2020; Szabadi, 2018). Electrical LC stimulation has been shown to evoke pupil dilation (Joshi et al., 2016; Liu et al., 2017) and the link between changes in pupil dilation, those not induced by luminance, and the LC-NE activity has been substantiated by a breadth of studies (Alnaes et al., 2014; De Gee et al., 2017; Gilzenrat et al., 2010; Murphy et al., 2011; Reimer et al., 2016; Varazzani et al., 2015), while correlations of pupil fluctuations with other neuromodulatory systems activity have also been described (Cazettes et al., 2021; Reimer et al., 2016). Pupil dilation has been proposed to scale with surprise and uncertainty about environmental statistics (Browning et al., 2015; Fan et al., 2023; Lawson et al., 2021; Nassar et al., 2012; Preuschoff, 2011).

To investigate pupil-linked neuromodulation during patch-leaving, we developed a novel version of a patch-foraging task for humans with two foraging environments varying in variability of patch initial rewards, which allowed us to quantify trial-wise pupil dilation in response to rewards preceding decisions to stay or leave. Based on MVT, we hypothesised adjustments of patch-leaving behaviour based on the instantaneous (‘foreground’) reward rate, thus shorter time spent in patches (‘patch residence times’) with small initial reward. Additionally, based on previous research showing that a patch-leaving decision variable is tracked by response times (RT, Kane et al., 2022), we expected that RT will be slower for leave than for stay decisions, while a continuous slowing down with an increasing patch residence times may be observed, presumably reflecting increasing decision difficulty with approaching the decision threshold. In line with previous research suggesting that increased LC-NE activity supports flexible adjustments of behaviour, reflected in pupil dilation, we hypothesised that leave decisions will be characterised by a larger transient pupil dilation than stay decisions. Similar to RT, this signal may increase across successive trials. We also anticipated that environmental variability may lead to higher levels of uncertainty (Kilpatrick et al., 2021; Nassar et al., 2012; A. J. Yu & Dayan, 2005), and therefore result in a sustained elevated pupil dilation in the environment with higher variability between patch values.

## Methods

### Participants

58 participants were recruited for the study at two testing locations: 24 at Ghent University (UGent) and 34 at Heinrich Heine University Düsseldorf (HHU). The recruitment took place via the SONA system and accompanying leaflets. Inclusion criteria were: normal or corrected-to-normal vision, no nystagmus, and an age between 18-30. Volunteer participants were reimbursed for their participation (20 EUR) or were given course credits. All participants were able to earn an additional bonus of maximum 10 EUR, determined based on their behavioural performance.

Two participants (HHU) were excluded due to incomplete data (not finishing the task) and one participant (UGent) was excluded due to technical failure with eye-tracking. Additionally, two participants (1 UGent, 1 HHU) were excluded based on the eye-tracking data exclusion criteria (see Data preprocessing). The final sample thus consisted of 53 participants (38 F and 15 M; *M*_age_ = 22.64, *SD*_age_ = 3.21).

The current study was approved by the ethical committee of the Faculty of Psychology and Educational Sciences of Ghent University (2021/147), and all participants provided written informed consent before taking part in the experiment.

### Stimuli and apparatus

Experimental stimuli were created and presented using Psychopy (Peirce et al., 2019). The stimuli were presented on a 24-inch computer monitor (resolution: 1440 × 900).

The stimuli consisted of: schematic drawings of bushes (either round or pointy depending on the environment and counterbalanced between participants) with berries (of 2 different colours for each environment counterbalanced between participants), a progress bar, two response icons (footprints and berries), and the travel time visualisation (clock counter and footprints), all presented on grey background. The luminance of the 7 colours that were used to create the stimuli and the background colour was matched as closely as possible for the particular testing location (*M_UGent_* = 5.92 cd/m^2^, SD = 0.27, range: 5.57–6.35; *M_HHU_* = 15.17 cd/m^2^, SD = 1.33, range: 13.40–17.00). This was done to minimise the effects of luminance on pupil size. The monitor was calibrated using a gamma correction procedure implemented in Psychopy.

### Patch – leaving task

In the patch-leaving task (see Fig. 1A), participants faced decisions between either continuing to collect rewards from a patch (represented as a blueberry bush) with exponentially diminishing rewards or moving (travelling) to another patch with replenished rewards, while foregoing reward collection during the travel time. To investigate the MVT – derived prediction that the time spent in the patch will vary depending on the initial patch reward rate, patches with three different initial rewards (small, medium, and high number of berries on the blueberry bush, see Fig. 1B) were presented. Additionally, two environments (farms) with differing levels of environmental variability in terms of reward distribution (high, HVAR, and low, LVAR) were included, signalled to participants by different shapes and colours of berries in each environment, to explore whether the variability of reward distribution in the environment influences patch-leaving and its pupillometric signature. The variability was manipulated through initial values of small and large patches, which were more extreme in the case of HVAR, while the average reward rate was kept close to identical in HVAR and LVAR (see Fig. 1B).

**Figure 1.**
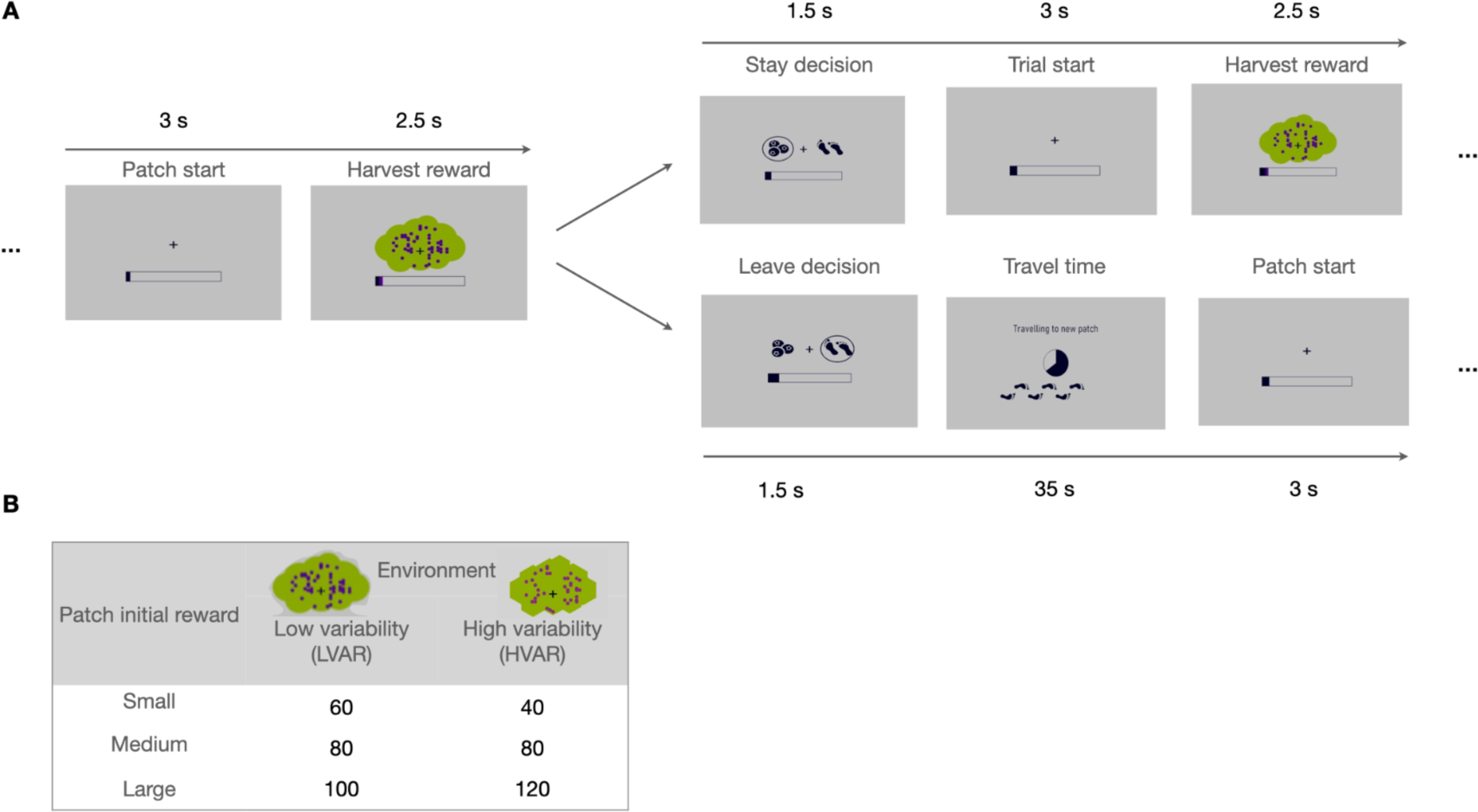
| Experimental paradigm. **A**: example of the initial trial in the patch and the transition in case of both stay decision and leave decision. Each trial starts with a centrally presented fixation cross for 3 seconds, after which a blueberry bush is presented for 2.5 seconds of passive harvesting. The participant is subsequently presented with the option to continue harvesting from the current bush (patch) or to move to a new bush. The options are represented with icons (berries and footprints for stay and leave decisions respectively), for a maximum of 1.5 seconds. Upon deciding to stay, the same trial sequence starts anew with a now decreased number of berries. Upon deciding to leave, the participant sees a travel screen for 35 seconds (during which no reward can be acquired), after which a new trial starts with a replenished bush. **B: Design of the task:** values of initial patch reward values in the high and low variability environment. Different environments were represented by different shapes and colours of bushes (counterbalanced between participants).

### Procedure

The experiment was conducted in an experimental room with dim and constant lighting. After signing the written informed consent, participants were seated in front of the computer screen with their head stabilised by a chin and forehead rest. Participants foraged for rewards by iteratively visiting the aforementioned virtual patches in two distinct environments. They were told that they would help a farmer collect blueberries from different bushes (patches) in two different farms (environments), in which bushes differ in colour and shape (see Fig. 1). Their goal was to collect as many blueberries as possible within an allocated time (40 minutes in each farm/environment). They were told they will be asked about the difference in the environments when they finish the task.

Each trial started with a fixation period (3 s), during which a fixation cross was presented centrally. Afterwards, a period of passive harvesting started with centrally-presented schematic drawing of a blueberry bush (2.5 s), together with a progress bar below the bush to signal the number of berries already collected. The number of randomly distributed circles on the bush represented the number of berries and thus the reward that was earned in this trial. The progress bar, which filled up several times over the course of the experiment, represented the number of berries already collected on previous trials (dark purple) and the current trial (light purple).

After this period of passive harvesting, two icons were presented to the left and right of the fixation cross, prompting participants to make a stay or leave decision (see Fig. 1). The icon symbolising berries was used to represent a stay decision, while the icon symbolising steps was used to represent a leave decision. The location (left or right of the fixation cross) of the icons was randomised on each trial in order to minimise motor preparation during the harvest time-window. Participants were asked to respond as quickly as possible by pressing a corresponding button (‘F’ for the icon on the left and ‘J’ for the icon on the right) on the keyboard (e.g., if the berry icon was on the left and steps icon on the right, the participant would press ‘F’ to stay, and ‘J’ to leave). The response window was limited to 1500 ms. If no response was provided within the time limit, a screen prompting participants to respond was presented (these responses were labelled as late, and were not considered in the analyses).

If participants indicated their decision to stay, the next trial started with a fixation period, and the amount of reward (the number of berries shown) in the subsequent harvest time window decreased exponentially (decay rate = 0.14). When participants made a decision to leave, a 35-s travel time started. During the travel time, a clock counter appeared in the center of the screen and was continually filled, while a series of moving footprints was shown in the lower visual field (see Fig. 1). After the travel time finished, a new, replenished patch was presented, in which the initial reward (i.e., number of berries) was drawn from one of the three values for this particular environment (see Table 1). The order of patches was pseudorandomised, thus in the series of 6 consecutive patches, each patch was present twice, in order to balance the overall reward rate between participants.

**Table 1.** | Behaviour in the foraging task: descriptive statistics. Mean (SD) of simulated optimal patch residence times, observed patch residence times, and overharvesting (results of one-sample Wilcoxon signed rank test against zero showing significant overharvesting in all conditions).

| Patch Type | Low variability (LVAR) |  |  | High variability (HVAR) |  |  |
| --- | --- | --- | --- | --- | --- | --- |
|  | Optimum Residence | Observed Residence | Overharvest | Optimum Residence | Observed Residence | Overharvest |
| Small | 4.75 (0.96) | 6.51 (2.98) | 1.76 (3.08)<br>V= 52626<br>$p < 0.001$ | 1.91 (1.01) | 4.58 (2.78) | 2.67 (2.87)<br>V= 73309<br>$p < 0.001$ |
| Medium | 6.69 (0.88) | 8.21 (3.38) | 1.51 (3.53)<br>V = 50022,<br>$p = 0.003$ | 6.42 (1.31) | 8.02 (3.26) | 1.60 (3.80)<br>V= 52574<br>$p < 0.001$ |
| Large | 8.70 (0.80) | 9.18 (3.27) | 0.48 (3.36)<br>V = 40346,<br>$p = 0.003$ | 9.08 (0.89) | 9.57(3.21) | 0.49 (3.53)<br>V= 40646<br>$p = 0.004$ |

The experimental blocks lasted approx. 20 minutes each, and participants performed 2 blocks in each environment, resulting in a total time of 40 min spent in the HVAR and LVAR environment, respectively. The order of the environments (HVAR and LVAR), as well as the assignment of colours and bush shapes to a particular environment, was counterbalanced between participants. HVAR and LVAR blocks were interspersed with each other so that the order of blocks was either LVAR, HVAR, LVAR, HVAR or HVAR, LVAR, HVAR, LVAR. Prior to the experimental blocks, two 5-minute practice blocks in each environment allowed participants to experience the distribution of rewards in each environment and familiarise themselves with the task.

Participants could receive a bonus of max. EUR 10 for their performance, which was determined by the ratio of the number of berries they collected and the maximum (optimal) number of berries (see *Optimum patch residence simulation* below). Each filled up progress bar in the course of the experiment represented 50 cents.

### Eye-tracking

Participants’ gaze and pupil size were recorded continuously with a sampling frequency of 1000 Hz using an infrared Eyelink 1000+ Desktop mount (SR Research Ltd., Ontario, Canada), while the head was stabilised by the SR Research Head Support. By default, the left eye was tracked for monocular recordings, except for 6/53 participants where the right eye was tracked due to technical issues. An eye-tracking calibration procedure was performed before each block.

### Behavioural analysis

Behavioural data were processed in R 4.2.2 (R Core Team, 2022), in conjunction with R Studio 2022.12.0. Trials from the last patch in block 2 and block 4 were excluded from analyses (as these blocks were interrupted after exactly 45 min spent in each environment so they did not always end with a leave decision and thus did not accurately capture patch-leaving behaviour). Trials with wrong key presses (i.e., keys other than ‘F’ and ‘J’; *M_wrong_* = 0.05 %, SD*_wrong_* = 0.12 %) and trials with responses that did not fall within 1.5 s from the onset of the response prompt were excluded from the analysis of response times (*M_late_* = 0.82 %, SD*_late_* = 1.73 %). Patch residence times (quantified as the count of trials/harvests spent in each patch), as well as trial-wise response times were used as dependent variables in linear mixed effect models (LME). Additionally, we quantified an overharvesting variable as the difference between the actual patch residence times and the participant-specific optimum patch residence times.

#### Optimum patch residence simulation

The optimum patch residence times were determined individually for each participant’s task structure by simulating the obtained rewards for each combination of possible patch residence times (with the maximum of patch residence time of 10) for each patch type and environment. The patch residence times for each patch type that maximised the possible reward were chosen separately for each environment. The resulting simulated optimum patch residence times (see Table 2) were similar to the simulation based on the MVT formula (Stephens & Krebs, 1986). Overharvesting was then quantified as the difference between optimum patch residence for each patch type and the actual patch residence.

**Table 2.**
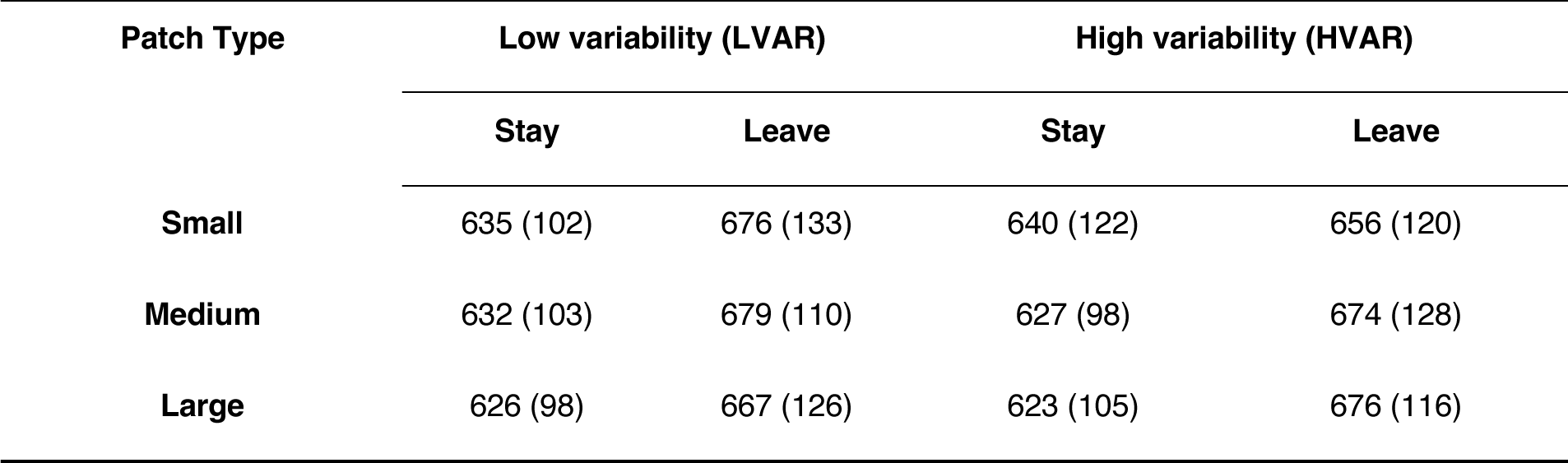
| RT and SD (in ms) split by Decision Type, Patch Type, and Environment Type.

| Patch Type | Low variability (LVAR) |  | High variability (HVAR) |  |
| --- | --- | --- | --- | --- |
|  | Stay | Leave | Stay | Leave |
| Small | 635 (102) | 676 (133) | 640 (122) | 656 (120) |
| Medium | 632 (103) | 679 (110) | 627 (98) | 674 (128) |
| Large | 626 (98) | 667 (126) | 623 (105) | 676 (116) |

#### Patch residence times analyses

For the analysis of patch residence times, we specified the factors environment (LVAR, HVAR), patch type (small, medium, large), previous patch type (small, medium, large; to quantify potential effects of transition between patches), patch number (mean-centred and scaled by the standard deviation; capturing the evolution of foraging behaviour with the time on task), and the interaction between environment and patch type as fixed effects in addition to a participant-specific random intercept:

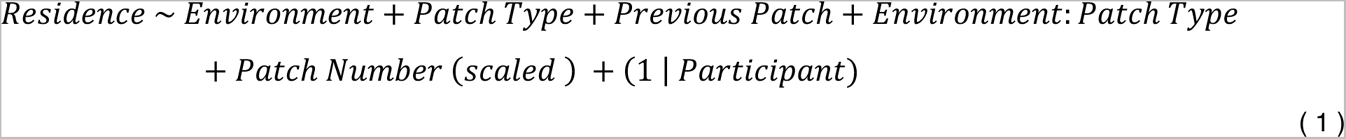

#### Overharvesting

The same model was fitted with overharvesting as the dependent variable:

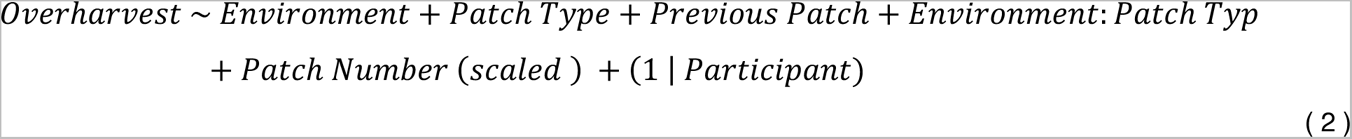

#### RT analyses

Trial-wise reaction times were modelled as a function of decision type (stay, leave), patch type (large, medium, small), and environment type (LVAR, HVAR) as fixed predictors, as well as a participant-specific random intercept:

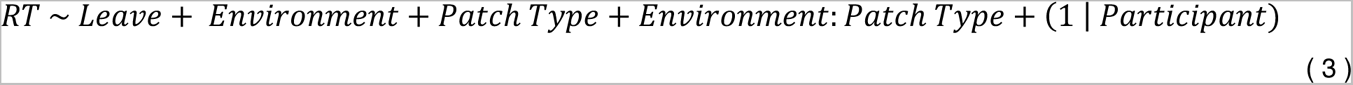

In order to further investigate the evolution of RT within patches, we isolated trials with stay decisions and assessed their linear dependence on Residence (mean-centred and scaled by the standard deviation), in a model that also included Environment type, Patch type, their interaction and a trial number (mean-centred and scaled by the standard deviation, to capture the time on task evolution):

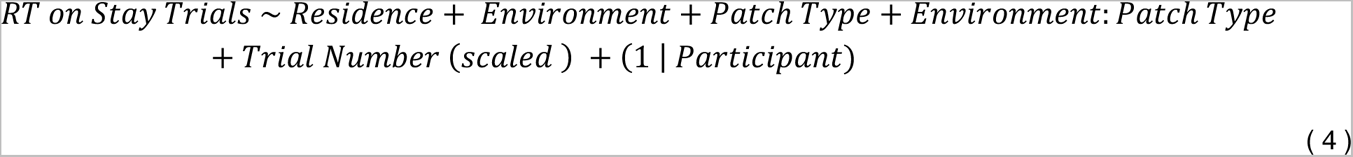

*F-* and *p*-values for fixed effects were estimated using the ‘lmerTest’ package (Kuznetsova et al., 2017), and the degrees of freedom were approximated using Satterthwaite’s method. Post-hoc contrasts were computed using estimated marginal means (EMMs) implemented in the ‘emmeans’ package (Lenth, 2017). Plots were generated using custom scripts and the ‘raincloudplots’ package (Allen et al., 2021).

### Pupillometry data preprocessing and analysis

#### Preprocessing

Eye-tracking data were preprocessed using Pupillometry Pipeliner (PuPl version 2.1.0; Kinley & Levy, 2021), installed on Matlab (The MathWorks Inc., 2021). Blinks were detected in continuous data using the pupillometry noise algorithm (Hershman et al., 2018). Residual small blinks were detected using the velocity profile method (Mathôt, 2013). Time points form 50 ms pre-to 150 ms post-blink were marked as missing data. Blinks were then linearly interpolated, with a maximum duration of interpolated segments of 800 ms, and the maximum deviation of interpolated data of 2 SD. Data were smoothed with a 150 ms mean Hann-windowed filter and segmented into epochs of –500 ms to 2500 ms around the onset of the evidence (bush and berries stimuli). Epochs with more than 33 % missing data were rejected from further analyses (*M_rejected_* = 2.47 %, *SD_rejected_* = 2.98 %). Data from participants with more than 25 % of epochs rejected before interpolation were excluded altogether. Two such participants were detected, one with 25.7 % of rejected epochs (UGent) and one with 38.9 % of rejected epochs (HHU). The mean number of trials included in pupillometry analyses was 390 (range: 251 – 519). Prior to analysis, data were baseline-corrected by subtracting the mean of 500 ms period preceding the onset of the stimuli display.

#### Time course analyses

In order to assess the temporal evolution of the task effects on pupil dilation, we conducted linear mixed effect models at each time point of the epoch (–0.5 – 2.5 s) with baseline-corrected pupil size values from each trial as dependent variables using the following model:

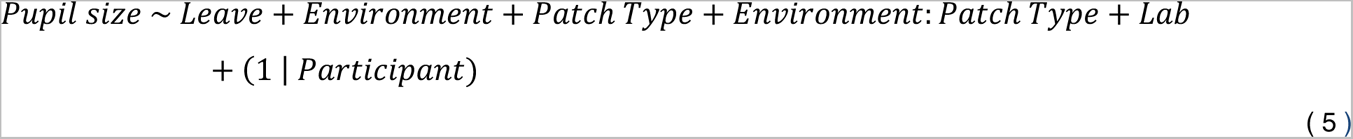

The resulting vectors of p-values were corrected for multiple comparisons using FDR correction (Benjamini & Hochberg, 1995).

#### Pupil dilation window (1 – 2 s) analyses

Further, we performed more detailed analyses on the averaged pupil size value from the interval of 1–2 s post-stimulus (bush presentation) onset. This window corresponded to the first maximum dilation of the pupil (following the initial constriction evoked by stimulus presentation), identified based on the average pupil trace across participants and conditions. We fitted the following model:

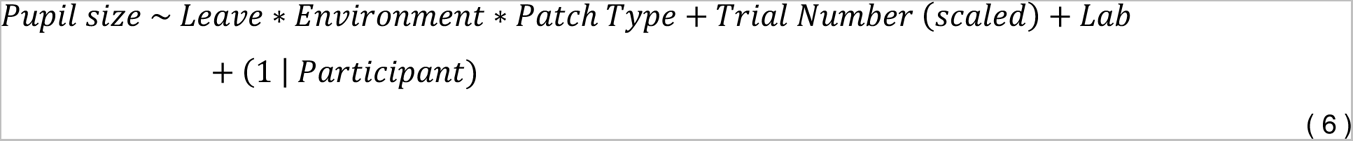

We also isolated pupil size in the stay decisions to assess whether it varies with patch residence times, patch type, and environment type, while controlling for trial number:

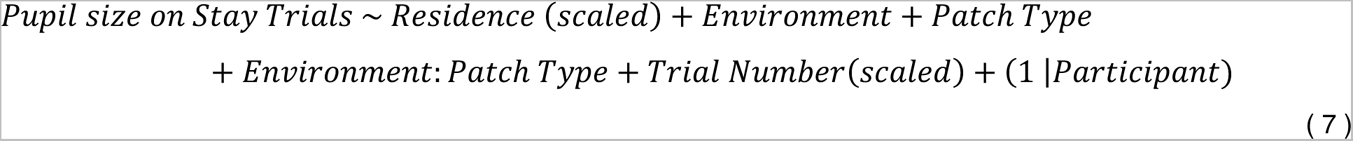

#### Behaviour and pupil size

To assess whether the evoked pupil response is related to behavioural effects, we ran three analyses. First, we ran a generalised linear mixed effects model estimating the log-odds of leave decisions based on the task variables and pupil dilation.

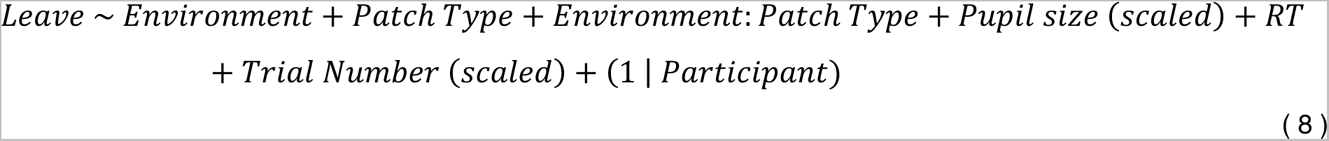

Second, we fitted an LME model predicting RT based on task variables and pupil dilation in the preselected time window (1–2 s), controlling for trial number.

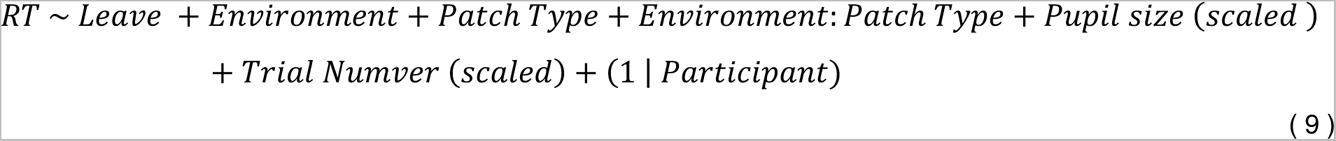

## Results

### Behaviour

#### Patch residence times analysis

The mean time spent in a patch was 7.64 (SD = 3.02, see Table 2 for descriptive statistics per experimental condition) trials. As predicted by MVT, patch residence times significantly varied depending on the patch type (Fig. 2A), *β_Medium–Small_* =3.26, 95 % CI (3.01, 3.50), *β_Large–Small_*= 4.78, 95 % CI (4.54, 5.03), *F*(2, 2664.9) 857.24, *p*<0.001, *η^2^_partial_ =* 0.39, 95 % CI (0.37, 1.00). Patch residence times were shorter in small reward patches than medium reward patches (estimate = –2.44, SE = 0.09, *t_ratio_* (2880) = –27.96, *p* < 0.001) and large reward patches (estimate = –3.74, SE = 0.09, *t_ratio_*(2880) = –42.84, *p* < 0.001, and also for medium vs large patches: estimate = –1.30, SE = 0.09, *t_ratio_*(2880) = –14.83, *p* < 0.001). There was a significant main effect of environment type, *β_LVAR–HVAR_* = 1.85, 95 % CI (1.61 2.10), *F*(1, 2664.8) = 51.47, *p*<0.001, *η^2^_partial_ =* 0.02, 95 % CI (0.01, 1.00), suggesting that, on average, participants harvested on average longer in the less variable environment. The interaction between patch type and environment type was also statistically significant (see Fig. 2A), *β_MediumPatch: LVAR_* = –1.71, 95 % CI (–2.06, –1.37), *β_LargePatch: LVAR_* = –2.30, 95 % CI (–2.64, –1.95), *F*(2, 2666) *=* 90.96, *p*<0.001, *η^2^_partial_ =* 0.06, 95 % CI (0.05, 1.00). Note that this interaction results naturally from the lower/higher patch values for the small/large patches in the HVAR compared to the LVAR environment. Post-hoc contrasts testing the effect of environment separately for each patch type showed that for large reward patches, patch residence times were significantly longer in the HVAR environment vs. LVAR environment, estimate = 0.44, SE = 0.12, *t*(2665) = 3.54, *p* < 0.001; whereas for small reward patches, patch residence times were significantly shorter for the HVAR vs. LVAR environment, estimate = –1.85, SE = 0.12, *t*(2665) = –14.82, *p* <0.001. For medium value patches, there was no significant difference between HVAR vs. LVAR environments, estimate = –.14, SE = 0.12, *t*(2665) = –1.14, *p* = 0.258. Patch residence times in the current patch were also dependent on previous patch type, *β_PrevMedium–PrevSmall_* =–0.03, 95 % CI (–0.21, 0.14),*β_Prev.Large arge–Prev.Small_*= –0.44, 95 % CI (–0.61, –0.26), *F*(2, 2664.8) *=* 14.81, *p* < 0.001, *η^2^_partial_ =* 0.01, 95 % CI (0.005, 1.00), being significantly longer when a previous patch contained small rewards, in comparison to when a previous patch contained large rewards, estimate = 0.44, SE = 0.09, *t*(2665) = 4.90, *p* < 0.001, and for previous medium reward patches compared to previous large reward patches, estimate = 0.40, SE = 0.09, *t*(2665) = 4.50, *p* < 0.001, while the comparison of previous small vs. previous medium reward was not significant, estimate = 0.03, SE = 0.09, *t*(2665) = .37, *p* = 0.927. A significant effect of patch number, *β_PatchNumber_* = –0.09, 95 % CI (–0.16, 0.001), *F*(2, 2670.20) *=* 4.89, *p* = 0.027, *η^2^_partial_ =* 0.002, 95 % CI (0.0001, 1), indicated that patch residence times generally decreased with the time on task.

**Figure 2.**
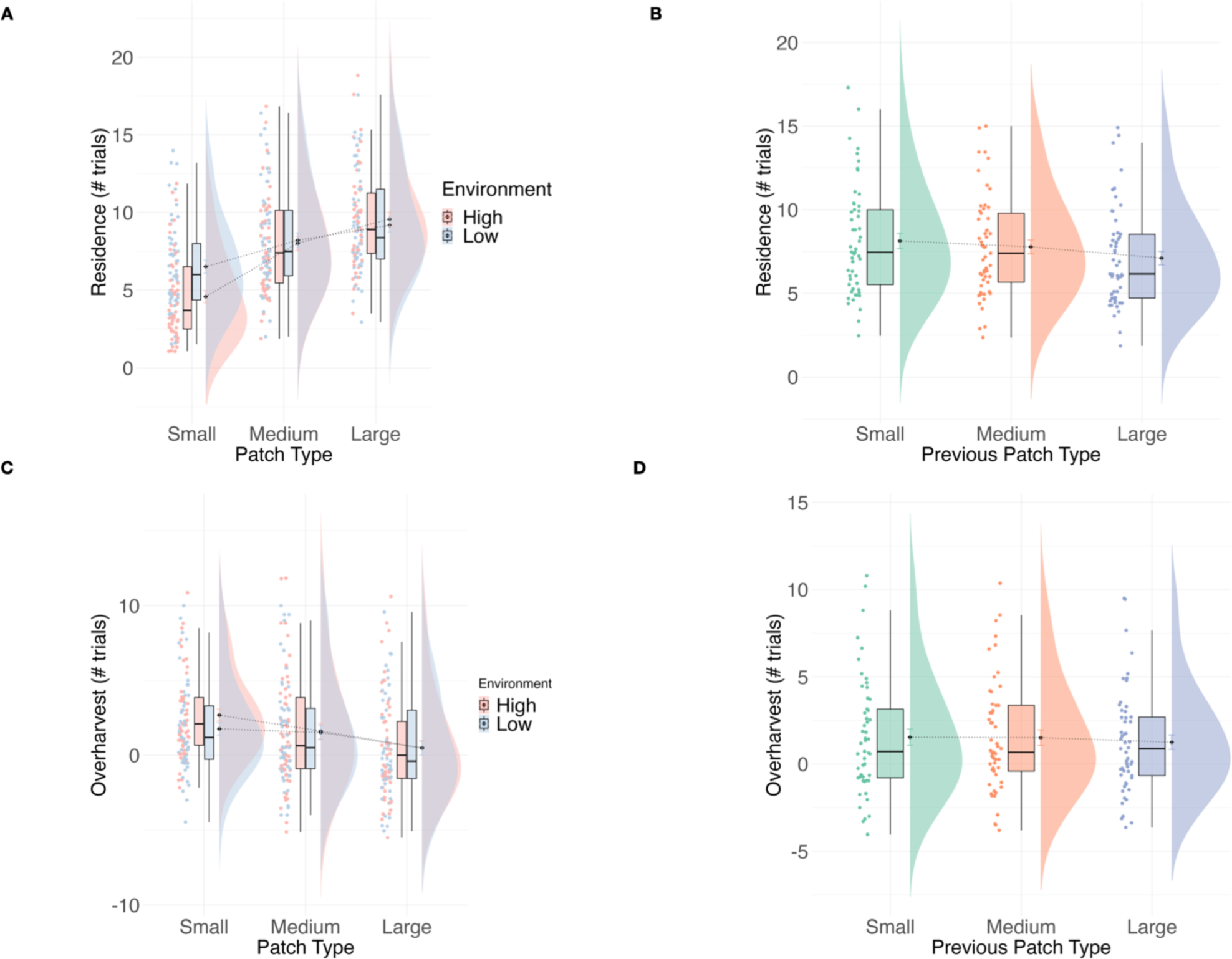
| Patch-leaving behaviour. **A**: Patch residence times patch type and environment type and **B:** previous patch type. **C:** Overharvesting split by patch type and environment type and **D:** previous patch type.

#### Overharvesting

Overall, participants stayed in patches longer overall than the optimum patch residence time (*M* = 1.41, *SD* = 3.13); see Table 1 for descriptive statistics per experimental condition. Overharvesting was most pronounced in the small patches and decreased with increasing patch value (Fig. 2B), as evident from a significant effect of patch type, *β_Medium–Small_* = –1.31, 95 % CI (–1.57, –1.04, *β_Large–Small_*= – 2.43, 95 % CI (–2.69, –2.16), *F*(2,2664.9) *=* 202.20, *p*<0.001, *η^2^_partial_ =* 0.13, 95 % CI (0.11, 1.00). All post-hoc comparisons between patches were significant, small vs. medium patches: estimate = 0.86, SE = 0.10, *t_ratio_*(2665) = 8.79, *p* < 0.001; small vs. large patches: estimate = 1.95, SE = 0.10, *t_ratio_*(2665) = 20.06, *p* < 0.001; medium vs. large patches: estimate = 1.09, SE = 0.10, *t_ratio_*(2665) = 11.17, *p* < 0.001. There was a significant main effect of environment type indicating more overharvesting in the high variability environment, *β_LVAR–HVAR_* = –0.98, 95 % CI (–1.24, –0.71), *F*(1, 2664.8) *=* 20.44, *p*<0.001, *η^2^_partial_ =* 0.008, 95 % CI (0.003, 1.00), and further, a significant interaction between patch type and environment type (see Fig. 2C), *β_LVAR:MediumPatch:_* = 0.90, 95 % CI (0.53, 1.28), *β_LVARt:LargePatch_* = 0.96, 95 % CI (0.58, 1.34), *F*(2, 2664.9) *=* 15.71, *p*<0.001, *η^2^partial =* 0.01, 95 % CI (0.005, 1.00), with post-hoc tests indicating that the overharvesting was larger for small patches in the HVAR environment compared to LVAR environment, estimate = 0.98, SE = 0.14, *t_ratio_*(2666) = 7.17, *p* < 0.001, while other comparisons did not reach statistical significance (all *ps* > 0.590). Further to this, the degree of overharvesting in the current patch also depended on the previous patch type (see Fig. 2D), *β_Medium–Small_* = –0.06, 95 % CI (–0.25, –0.13), *β_Large–Small_*= – 0.48, 95 % CI (–0.67 – 0.29), *F*(2,2664.8) *=* 14.26, *p* < 0.001, *η^2^_partial_ =* 0.01, 95 % CI (0.005, 1.00), with overharvesting being significantly larger when the previous patch contained small rewards compared to large reward patches, estimate = 0.48, SE = 0.10, *t_ratio_*(2666) = 4.90, *p* < 0.001, and when the previous patch contained medium rewards compared to large rewards, estimate = 0.41, SE = 0.10, *t_ratio_*(2666) = 4.25, *p* < 0.001. The previous small reward patch vs. previous medium reward patch comparison was not statistically significant, estimate = 0.06, SE = 0.10, *t_ratio_*(2666) = 0.63, *p* = 0.805. The effect of patch number did not reach significance, *β_PatchNumber_* = –0.05, 95 % CI (–0.14, 0.03), *F*(2, 2670.7) *=* 1.70, *p* = 0.192, *η^2^_partial_ =* 0.0006, 95 % CI (0.00, 1), indicating that the degree of overharvesting did not significantly vary with time on task.

#### Response times (RT) analysis

Participants’ overall mean RT was 634 (*SD* = 98) ms (see Table 2 for descriptive statistics per experimental condition). RTs for leave decisions (*M* = 671, *SD =* 106) were significantly slower than for stay (*M* = 628, *SD =* 97) decisions, *β_Leave–Stay_* = 42.62, 95 % CI (36.38, 48.86), *F*(1,20237) *=* 179.09, *p*<0.001, *η^2^_partial_ =* 0.008, 95 % CI (0.006, 1.00). Additionally, RT were slower in the LVAR (*M_LVAR_* = 637, SD = 99 ms) than the HVAR (*M* = 632*_HVAR_*, SD = 97 ms) environment, *β_LVAR–HVAR_* = 7.62, 95 % CI (–1.33, 16.57), *F*(1,20226) *=* 6.14, *p* = 0.013, *η^2^_partial_ =* 0.0003, 95 % CI (0.00003, 1.00).^1^

The analysis including stay trials only (see Behaviour Analysis, Eq. 4) indicated that across successive stay decisions leading up to a leave decision, response times gradually increased (see Fig. 3B), *β_Residence-scaled_* = 14.03, 95 % CI (11.58, 16.48), *F*(1,20273) = 126.11, *p* < 0.001, *η^2^_partial_* = 0.006, 95 % CI (0.004, 1). Furthermore, there was a significant main effect of patch type, *β_Medium-Small_* = –10.47, 95 % CI (–19.00, –1.94), *β_Large-Small_* = –17.14, 95 % CI (–25.52, –8.75), *F*(2, 20231) *=*15.38, *p* < 0.001, *η^2^_partial_ =* 0.001, 95 % CI (0.0007, 1.00). The average RT was slower in the small reward patches compared to the medium reward patches, estimate *=* 7.90, SE = 2.95, *t*(20230) = 2.68, *p* = 0.024, and large reward patches, estimate *=* 15.89, SE = 2.91, *t*(20236) = 5.45, *p* < 0.001 (medium vs large patches: estimate = 7.99, SE = 2.51, *t*(20228) = 3.19, *p* = 0.004). RT generally decreased with the time on task, as indicated by a significant main effect of trial number, *β_TrialNumber-scaled_* = –15.19, 95 % CI (–17.39, –13.00), *F* (1,20242) = 184.06, *p* < 0.001, *η^2^_partial_* = 0.009, 95 % CI (0.007, 1). The effect of environment type or the interaction between environment type and patch type were not significant, *p* > 0.178.^2^

**Figure 3.**
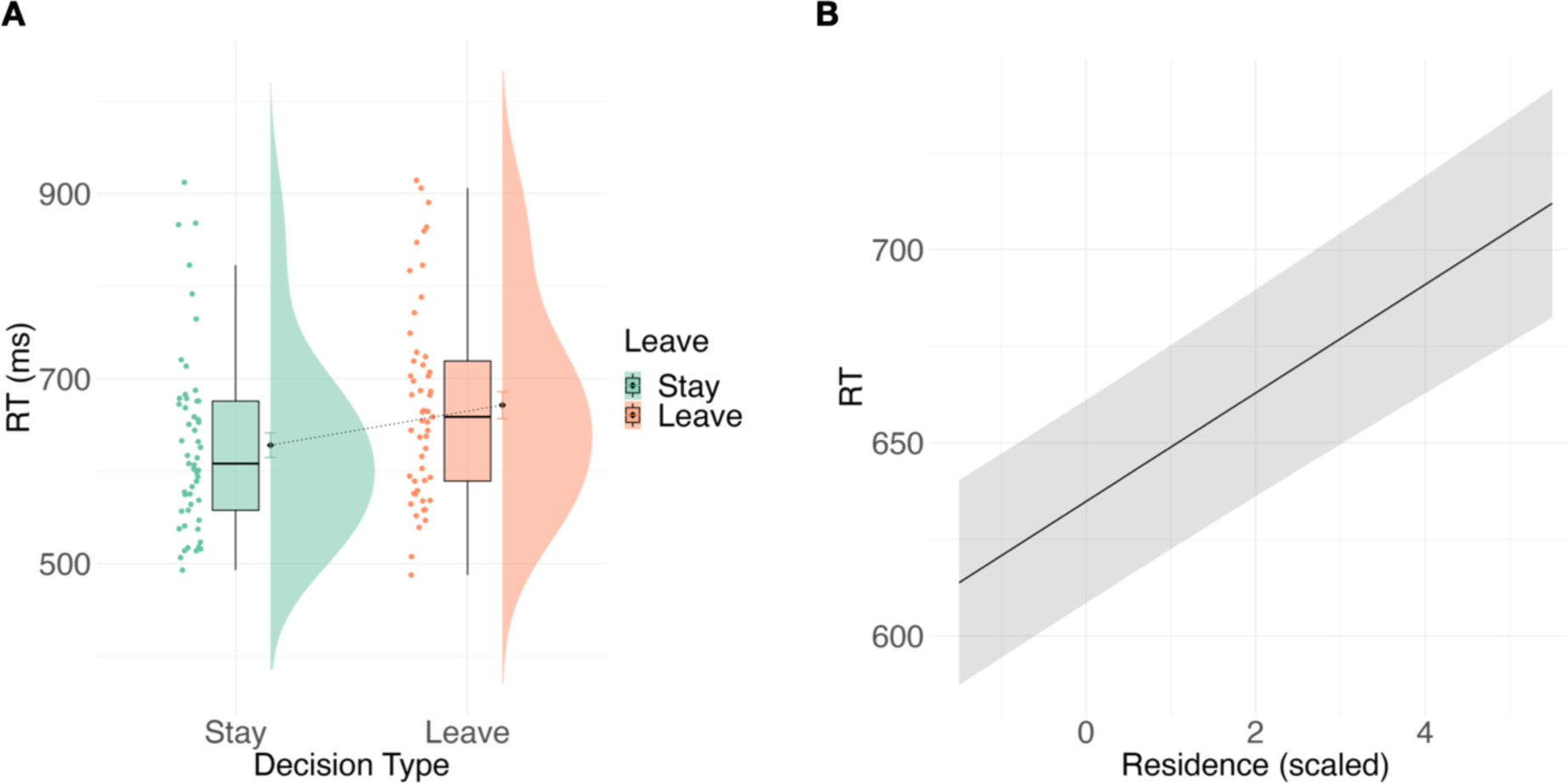
| RT effects. **A**: The effect of decision type on RT: RT slower for leave vs. stay decisions. **B:** The predicted RT on stay trial as a function of patch residence times: RT increases with successive stay decisions leading up to a leave decision.

### Pupillometry

#### Time course (–0.5 – 2.3 s) analyses

Following evidence (bush) presentation, the pupil response was characterised by a rapid and transient constriction, followed by a first dilation, peaking approx. around 1.5 s after the stimulus onset. Later, a more sustained dilation followed, increasing towards the onset of the decision window. The analysis performed on the entire time course (see Eq. 5) indicated that the pupil size was larger on trials with leave decisions than on trials with stay decisions (see Fig. 4A). This difference was significant between 467 ms – 2500 ms (FDR-corrected p-values <0.05), encompassing both the constriction and the dilation. This effect gradually increased over the course of the epoch (see Fig. 4D). Pupil size also differed as a function of patch type (see Fig. 4B), as indicated by a significant effect observed between 843 ms – 1243 ms (see Fig. 4D). Additionally, the evoked pupil size differed between testing locations, with values being higher for HHU than UGent participants (FDR-corrected *p*-values <0.05 between 845–1486 ms), an effect that may be attributed to differences in the ambient luminance. A main effect of the environment, and an interaction between patch type and environment type did not reach the significance level (all FDR-corrected *p*s > 0.05).

**Figure 4.**
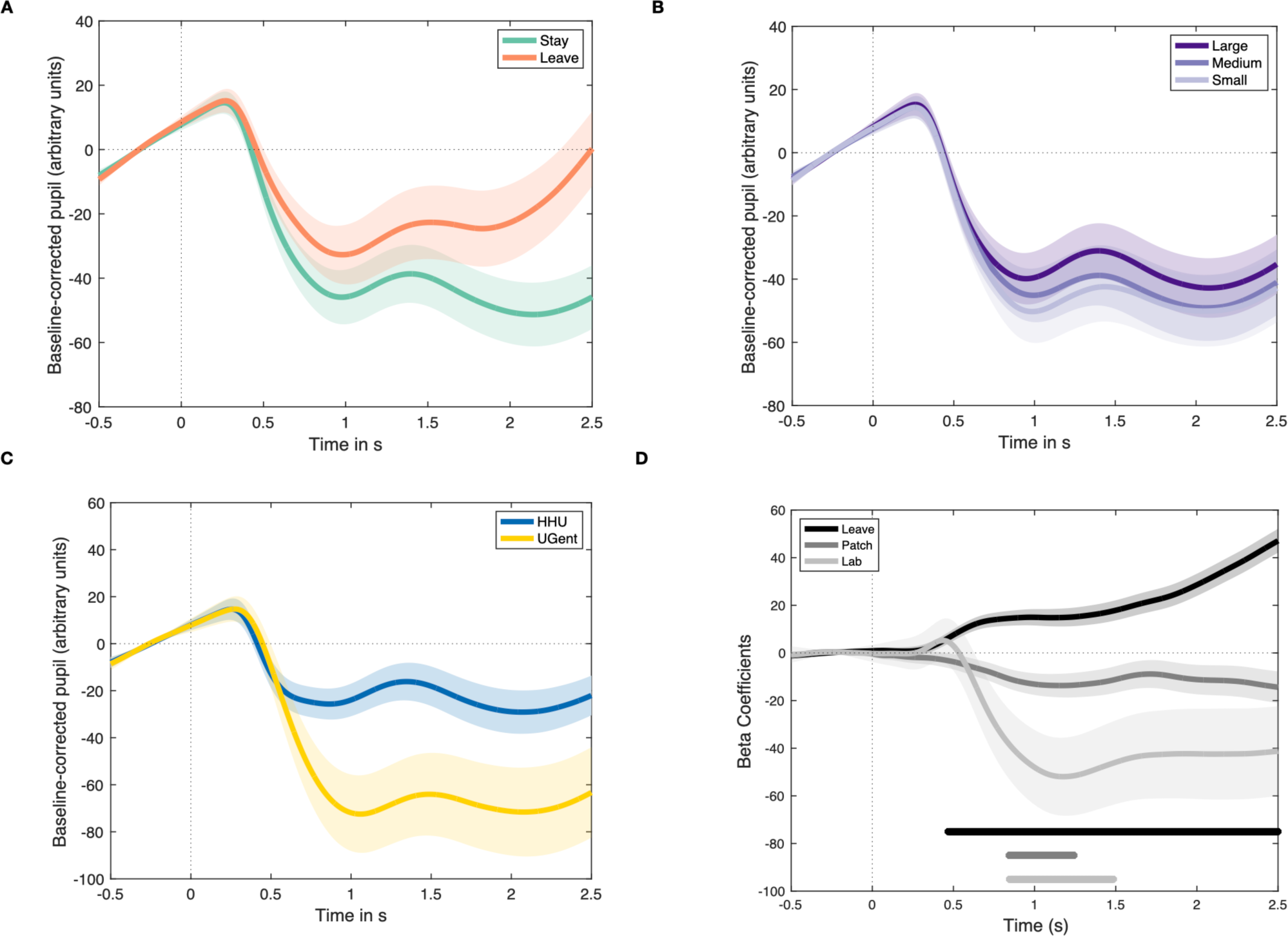
| Pupil size time series analyses. **A**: The effect of decision type on pupil size: Pupil size larger for leave vs. stay decisions. **B:** The effect of patch type on pupil size: Pupil size larger in the high vs. low variability environment. **C:** The effect of lab on pupil size: Pupil size larger at HHU vs. UGent. **D:** Beta coefficients for the effects of decision type, patch type, and lab derived from LME analyses (Eq. 5) on the pupil time series. Horizontal lines indicate significant FDR-corrected p-values.

#### The pupil dilation window analysis (1 – 2 s)

The analysis in the time window corresponding to the first dilation of the pupil after the evidence onset (see Eq. 6) showed a significantly larger average dilation on trials with leave decisions compared to stay decisions (see Fig. 5A), *β_Leave– Stay_* = 49.35, 95 % CI (29.52, 69.17), *F*(1,19782.9) *=* 22.69, *p*<0.001, *η^2^_partial_ =* 0.001, 95 % CI (0.0005 1.00), in line with the above analyses of the entire time course. Furthermore, there was an effect of the environment on the pupil size (Fig 5B). Average pupil dilation was significantly larger in the HVAR than in the LVAR environment, *β_LVAR–HVAR_* = –1.93, 95 % CI (–14.67, 10.81), *F*(1,19747.9) *=* 15.18, *p*<0.001, *η^2^_partial_ =* 0.0008, 95 % CI (0.0003, 1.00). The main effect of patch type on pupil dilation was not significant, *β_Medium–Small Patch_* = 14.94, 95 % CI (2.91, 26.98), *β_Large– Small Patch_*= 19.64, 95 % CI (7.97, 31.30), *F*(2, 19749.4) *=* 0.90, *p* = 0.405, *η^2^_partial_ =* 0.00009, 95 % CI (0.00, 1.00), however, there was a significant relationship between decision type and patch type, showing that on stay trials, pupil size was significantly smaller in the small compared to large reward patches, estimate = –16.19, SE = 3.99, *t*(19766) = –4.06, *p* < 0.001, while the difference between medium and large reward patches, estimate = –7.94, SE = 3.39, *t*(19753) = –2.34, *p* = 0.051, and small vs. medium reward patches, estimate = –8.25, SE = 4.10, *t*(19755) = –2.01, *p* = 0.110 did not reach significance. Nor were the comparisons of patch types on leave trials significant, *p* > 0.177. Pupil size also increased with increasing trial number, *β_Trialnumber-scaled_ =* 5.82, 95 % CI (3.05, 8.59), *F*(1, 19792.7) *=* 16.92, *p* < 0.001, *η^2^_partial_ =* 0.0008, 95 % CI (0.0003, 1.00), while also differed as a function of testing location, being larger at HHU than at UGent, *β_UGent – HHU_ =* –46.36, 95 % CI (–80.41, –12.32), *F*(1,50.9) = 7.12, *p* = 0.010, *η^2^_partial_ =* 0.12, 95 % CI (0.02, 1.00). All other interactions were not significant, all *p* > 0.089.

**Figure 5.**
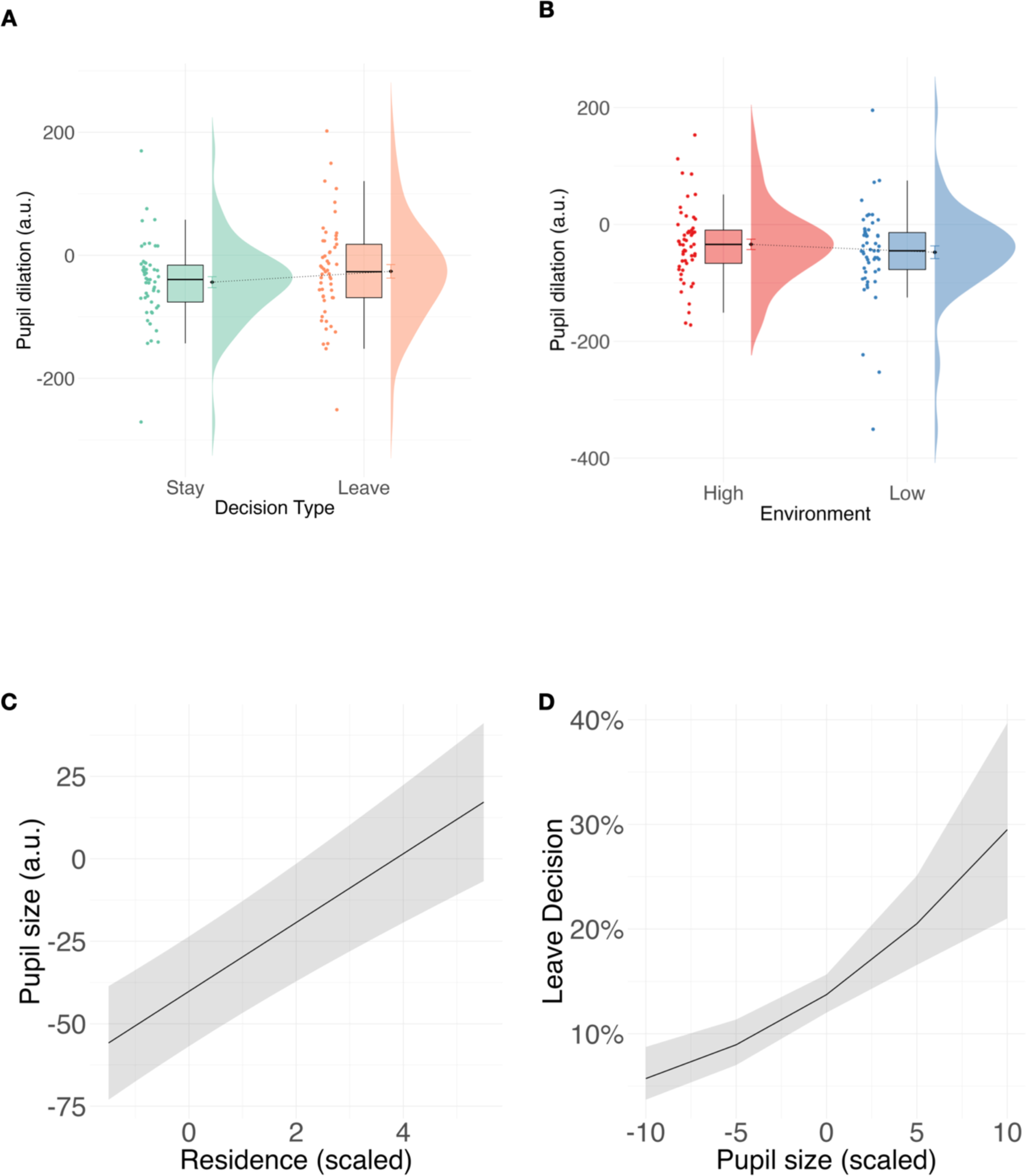
| Pupil size values in the window of the first dilation (1–2 s): **A**: The effect of decision type: a greater dilation for leave vs. stay decisions. **B:** The effect of environment: larger dilation in the high vs. low variability environment. **C:** Pupil dilation increases with the time spent in the patch on stay trials. **D:** Pupil size significantly predicts the odds of leave decisions.

The average pupil dilation on stay trials (see Eq. 7) significantly increased with the time spent in the patch, *β_Residence-scaled_* = 10.43, 95 % CI (7.34, 13.52), *F*(1,19623.4) = 43.65, *p* < 0.001, *η^2^_partial_* = 0.002, 95 % CI (0.001, 1). Pupil size also significantly varied as a function of environment type, being larger in the HVAR vs LVAR environment, *β_LVAR–HVAR_* = –13.22, 95 % CI (–24.51, 1.92), *F*(1,19756.6) *=* 21.02, *p* < 0.001, *η^2^_partial_ =* 0.001, 95 % CI (0.0004, 1.00). Finally pupil size significantly increased with increasing trial number, *β_Trialnumber-scaled_ =* 5.93, 95 % CI (3.16, 8.71), *F*(1, 19797.7) *=* 17.62, *p* < 0.001, *η^2^_partial_ =* 0.0009, 95 % CI (0.0003, 1.00). Pupil size was smaller at UGent than at HHU, *β_UGent – HHU_ =* –45.61 95 % CI (–79.36, –11.86), *F*(1,50.9) = 7.02, *p* = 0.011, *η^2^_partial_ =* 0.12, 95 % CI (0.02, 1.00). The effect of patch type or the interaction between patch type and environmental variability was not significant, *p* > 0.221.

#### Pupil and behaviour

Pupil dilation was predictive of patch-leaving decisions. The GLMM model (Eq. 9) indicated that pupil dilation significantly predicted the odds of making a leave decision, with larger pupil sizes being associated with an increased probability of leaving a patch, *β_Pupil_* =0.10, χ^2^ = 19.55, *p* < 0.001, when controlling for other variables. In contrast, RT was not significantly predicted by the average pupil dilation in the interval from 1 – 2 s after the onset of the evidence (Eq. 9), *β_Pupil dilation_* = 0.008, 95 % CI (–0.005, 0.02), *F*(1,19777) = 1.44, *p* = 0.229, *η^2^_partial_* = 0.00007, 95 % CI (0.00, 1).

## Discussion

We investigated pupil dilation responses as a non-invasive index of brainstem neuromodulation in a patch-leaving task. Based on previous research on the possible role of the LC-NE system in the control of patch-leaving decisions, we hypothesised that the decisions to abandon the current resources in search for new alternatives would be associated with a transient noradrenergic activation, which would be reflected in an increased pupil dilation. Our results showed a clear pupillometric signature of patch-leaving: decisions to leave the patch were associated with the trial-by-trial evoked pupil dilation following the presentation of the evidence (reward), prior to the response window. Pupil dilation was larger for leave than for stay decisions, and this difference emerged rapidly (∼450 ms) after the reward presentation. The effect of decision type ramped up throughout the epoch, towards the onset of the response window. Furthermore, across successive stay decisions leading up to a leave choice, a continuous increase in pupil dilation was observed.

Behaviour across two testing locations principally conformed with predictions derived from the MVT (Charnov, 1976; Stephens & Krebs, 1986): participants adjusted their patch residence times to the foreground reward value of patches, and left the patches with small initial reward values earlier than those with larger initial rewards, corroborating previous studies (Kane et al., 2019; Le Heron et al., 2020). However, a tendency to overharvest, a phenomenon frequently described in studies across species (Doren et al., 2023; Harhen & Bornstein, 2023; Kane et al., 2019; Kendall & Wikenheiser, 2022), was also present in the current study. Further, in contrast with the MVT prediction that all patches would be abandoned at the same threshold equal to the average reward rate, the degree of overharvesting varied depending on the initial reward rate, being larger in patches with smaller initial reward value. Furthermore, a longer-than-optimal residence in the environment with higher variability of reward values was observed, due to the larger degree of overharvesting in the patch type with the initial smallest rewards. While some previous studies reported a uniform degree of overharvesting across patch types (Kane et al., 2019; Le Heron et al., 2020), a recent study reported a larger degree of overharvesting for low-value options (Harhen & Bornstein, 2023; for a study in rats, see: Kane et al., 2022; note an opposite pattern: Doren et al., 2023). Such a pattern was shown to emerge computationally due to uncertainty about patch reward distributions or poor resource availability (Kilpatrick et al., 2021). Additionally, current results suggest that overharvesting may be brought about by the transition structure in the environment (Harhen & Bornstein, 2023; Yoon et al., 2018), with an increased tendency to overharvest upon transitioning from previously encountering smaller rewards.

Participants’ RTs were in general slower for leave decisions, moreover, they continually increased with longer patch residence times, and thus, with proximity to leave decisions. Such a pattern, corroborating previous findings from rodents (Kane et al., 2022), has been thought to reflect an incrementally increasing decision difficulty as the threshold to leave is approached (Kane et al., 2022; Shenhav et al., 2014). Meanwhile, an overall decrease of RT observed with the time on task may be attributed to familiarisation with the task structure (Willemet, 2024) and possibly, a reduction of decision uncertainty (Urai et al., 2017). In turn, behavioural invigoration by high expected reward value (Willemet, 2024; Yoon et al., 2018) could account for faster RTs for larger compared to smaller reward patches.

Consistent with our prediction that pupil fluctuations may track environmental variability, we observed that pupil size in the dilation time window was overall larger in the environment with higher variability between patches. Of note, this effect did not survive multiple comparison correction in the entire time course analysis of pupil size, but it was significant in the dilation time window, as well as in the analysis of the derivative of pupil time series (see Supplementary Material 3. *Time course analysis of pupil derivative*). As pupil dilation is believed to track uncertainty and environmental volatility estimates (Browning et al., 2015; Lawson et al., 2021; Nassar et al., 2012), tracking of the increased variability of the rewards distribution may have resulted in an increased pupil dilation. How different types of uncertainty influence pupil dilation is a matter of ongoing research (Filipowicz et al., 2020; Grujic et al., 2024; Nassar et al., 2012). Variability of the environmental statistics, or estimation uncertainty, can be computationally expressed by estimates of ‘expected uncertainty’ (A. J. Yu & Dayan, 2005) or ‘relative uncertainty’ (Nassar et al., 2012). Relative uncertainty estimates were previously linked to increases in mean baseline pupil size (Nassar et al., 2012; note also an effect of total uncertainty in Fan et al., 2023). Our supplementary analyses also point to the elevated baseline pupil values with the higher environmental variability (See Supplementary Material 2. *Controlling for baseline effects in pupillometry data*). Increased tonic LC activity, which may be reflected in elevated baseline pupil values (Aston-Jones & Cohen, 2005; Gilzenrat et al., 2010; Jepma & Nieuwenhuis, 2011), has been shown to lead to earlier patch leaving brought about by an increased decision noise in rats (Kane et al., 2017), and to a higher probability of behavioural switches in mice (McBurney-Lin et al., 2022). Pharmacologically decreasing noradrenaline release also resulted in a decrease in choice variability during an operant effort task in monkeys (Jahn et al., 2018).

In turn, elevated baseline pupil values were linked to a disengagement, facilitating exploratory choices (Jepma & Nieuwenhuis, 2011), and particularly to uncertainty-driven random exploration (Fan et al., 2023). The current study revealed differences in patch-leaving behaviour due to environmental variability, such as slightly faster RTs in the higher variability environment, which could indicate effort mobilisation (Jahn et al., 2018) and a higher degree of overharvesting (specifically in the small value patch), a pattern that may emerge as a consequence of uncertainty (Kilpatrick et al., 2021) or variability in choices (Cash-Padgett & Hayden, 2020). Furthermore, current results indicate that disengagement from current options in a patch-leaving task is tracked by increases in transient event-related, rather than baseline (‘tonic’) pupil size measurements.

Our results provide the first, to our knowledge, direct evidence that in a patch-foraging scenario, an increased event-related (‘phasic’) pupil-linked arousal increases the odds of abandoning current diminishing resources for alternative options. Previous findings show that elevated phasic pupil dilation encodes surprise (Alamia et al., 2019; O’Reilly et al., 2013; Preuschoff, 2011; Zhao et al., 2019), information gain (Zénon, 2019), and is related to task-switches (Da Silva Castanheira et al., 2021; Katidioti et al., 2014; Rondeel et al., 2015), choice alterations (Urai et al., 2017), decrease of perceptual and choice biases across species and tasks (De Gee et al., 2017, 2020). All of these effects might translate into an increased behavioural variability that could manifest as an increased drive to switch and explore alternative options (Grujic et al., 2024).

The pupillometric signature of patch-leaving observed in the current study seems consistent with the network reset theory of LC activity, which proposes that phasic LC bursts reset ongoing activity in cortical networks, thereby enabling switching of behavioural states in order to flexibly adapt to environmental contingencies (Bouret & Sara, 2005; Grella et al., 2019; Karlsson et al., 2012; Sales et al., 2019; Sara & Bouret, 2012). On the other hand, based on the adaptive gain theory (Aston-Jones & Cohen, 2005), it could be predicted that increased tonic LC activity drives disengagement from current options in search for more valuable alternatives (Fan et al., 2023; Jepma & Nieuwenhuis, 2011), which is not what we observed. Conversely, lower pre-stimulus baseline pupil values were observed preceding leave decisions. Crucially, the pattern of event-related increase in pupil size preceding leave decisions remained significant when data were analysed with no baseline correction applied, including trial-by-trial baseline values as a regressor (cf. Alday, 2019; see Supplementary Material 2. *Controlling for baseline effects in pupillometry data*). Additional analyses with the derivative of pupil time series, proposed to be less susceptible to baseline effects (Filipowicz et al., 2020) and a more accurate predictor of cortical states (Reimer et al., 2016; Yang et al., 2021), revealed a consistent patch-leaving effect, although clearly present in two separate time windows (early window comprising constriction and dilation and a late windo*w* with a ramping dilation, see Fig. S2). Both analyses convergently revealed a prolonged ramping up of pupil dilation towards the decision window, a pattern which is comparable to the effect of tonic electrical LC stimulation on pupil size (Liu et al., 2017), and may thus indicate a tonic mode effect.

The extent to which ‘phasic’ and ‘tonic’ pupil dilation responses map onto phasic and tonic modes of LC-NE activity is a matter of current debate (Grujic et al., 2024; Joshi & Gold, 2020). While some previous studies have assumed a relatively direct mapping between baseline pupil and the tonic, and task-evoked dilation and the phasic mode of LC firing (Cole et al., 2022; Gilzenrat et al., 2010), recent findings qualify this link by emphasising considerable variability and a relatively low temporal precision of LC – pupil coupling (Joshi & Gold, 2022; Megemont et al., 2022; Yang et al., 2021). Other neuromodulatory systems were also noted to be co-active with pupil diameter fluctuations (Larsen & Waters, 2018; Reimer et al., 2016). Basal forebrain acetylcholinergic (ACh) activity was shown to co-vary with pupil size (Nelson & Mooney, 2016; Reimer et al., 2016), specifically at longer time-scales than NE – pupil correlation, consistent with the theoretical notion that baseline pupil fluctuations may index expected uncertainty tracked by ACh (Marzecová et al., 2019; Nassar et al., 2012; A. J. Yu & Dayan, 2005). Additionally, pupil dilation has been related to brainstem serotonergic (Cazettes et al., 2021; Y. Yu et al., 2004), and even LC-GABAergic influences (Breton-Provencher & Sur, 2019).

We propose that the event-related increase of pupil dilation related to leave decisions may be indicative of an increased LC-NE activity. Nevertheless, complex patch-leaving behaviour is orchestrated by co-activation of several other neuromodulatory systems. Dopaminergic activity is thought to modulate the perceived opportunity cost of time (Constantino et al., 2017; Ianni et al., 2023; Le Heron et al., 2020). In contrast to sustained LC-NE stimulation causing earlier patch-leaving (Kane et al., 2017), a sustained DR stimulation increased persistence, thus leading to later patch-leaving times (Lottem et al., 2018), suggesting a link of serotonergic activity with exploitation states (Marques et al., 2020). Administration of nicotine – an ACh agonist – decreased overharvesting in humans, leading to more optimal performance (Doren et al., 2023), while interestingly, the same study reported a less pronounced decrease in patch residence times under reboxetine – a NE reuptake inhibitor. Psychopharmacological NE manipulation employed by several previous studies likely leads to sustained shifts in both tonic and phasic NE levels (Cremer et al., 2022; Hauser et al., 2018; Lawson et al., 2021), and the temporal fluctuations of LC-NE firing related to patch-leaving remain to be further investigated, while pupillometry can be utilised to index the temporal fluctuations in brainstem arousal states.

Convergent evidence from neuroimaging and electrophysiological studies suggests the role of the dorsal anterior cingulate cortex (dACC) activity in controlling patch-leaving decisions (Hayden et al., 2011; Kaiser et al., 2021; Karlsson et al., 2012; Kolling et al., 2012). Causal evidence implicates ACC microcircuitry in arbitrating between persisting with ongoing behavioural strategy or switching away (Tervo et al., 2021), while LC input to ACC has been shown to govern a transition to a stochastic mode of behaviour (Tervo et al., 2014). Ramping of dACC neuron firing with increasing patch residence times has been observed, possibly indicative of encoding a decision threshold (Hayden et al., 2011; Kane et al., 2022). This ramping is reminiscent of our pattern of pupil dilation linked to patch-leaving decisions, a possible relationship that is supported by findings showing co-variation between ACC activity and pupil changes (Joshi et al., 2016, Ebitz & Platt, 2015). Furthermore, the increase in dACC firing rate with patch residence time was also reported to scale with RT (Kane et al., 2022). Despite a similarity in the pattern of ramping event-related pupil dilation and RT towards leave decisions in the current study, we did not observe a significant trial-by-trial correlation between RT in the decision window and the pupil size in the early dilation window, suggesting that early pupil dilation and RT effects may reflect partially distinct subprocesses in the current patch-leaving context.

A factor limiting our descriptions of the pupil dynamics linked to patch-leaving was the observation of a continuous decrease of baseline pupil values with the time spent in the patch, and the time on task (see Supplementary Material 2 *Controlling for baseline effects in pupillometry data*), previously described in the literature (Fan et al., 2023; Skora et al., 2024; Unsworth et al., 2019; Unsworth & Robison, 2016). Our analyses suggest that the patch-leaving effect, although partially affected by the temporal drift, remains significant when controlling for the decreasing baseline values with the time on task. Such decreases were attributed to vigilance decrements and fatigue (Hopstaken et al., 2015; Martin et al., 2022; McLaughlin et al., 2023). However, this interpretation does not seem to concur with the observed speeding up of RT with the time on task, and with the theoretical notion that decreasing baseline pupil values may be associated with a transition to phasic LC mode (Jepma & Nieuwenhuis, 2011), which is thought to increase attentional engagement (Aston-Jones & Cohen, 2005). In line with this notion, a decrease in patch residence times with the time on task has been observed previously (Barack et al., 2023, 2024; Harhen & Bornstein, 2023), as well as in the current study. Note that in our study, it did not seem to translate into a decreased overharvesting tendency. Additionally, the pupillometric signature of patch leaving remains present when controlling for the effect of testing location, resulting from different levels of baseline pupil values (see Supplementary Material 2 *Controlling for baseline effects in pupillometry data*), likely related to difference in illuminance at the two testing locations due to ambient lightning (Mathôt & Vilotijević, 2022), while not having a detectable effect on behaviour (See Supplementary Material 1 *Controlling for the effects of testing location on behaviour*).

In conclusion, we propose that patch-leaving decisions are associated with increased brain-stem arousal as reflected in event-related pupil dilation, presumably indicating an increased LC-NE activity associated with abandoning current options and prompting behavioural adjustments. The pupillometric signature of patch-leaving may be utilised in future studies describing neuromodulatory dynamics of foraging behaviour.

## Acknowledgements

We thank Monja Froböse for methodological and theoretical advice. This research was supported by the Deutsche Forschungsgemeinschaft (DFG) grant (JO-787/6-1) and by the Research Foundation Flanders (FWO) postdoctoral fellowship (12V5620N).

## CRediT authorship contribution statement

**Anna Marzecová:** Conceptualisation, Methodology, Software, Validation, Formal analysis, Investigation, Resources, Data Curation, Writing – Original Draft, Writing – Review & Editing, Visualisation, Supervision, Project administration, Funding acquisition

**Brent Vernaillen:** Conceptualisation, Methodology, Software, Formal analysis, Investigation, Resources, Data Curation, Writing – Review & Editing, Visualisation

**Ida Hoxhaj:** Formal analysis, Investigation, Data Curation, Writing – Review & Editing

**Luca F. Kaiser:** Conceptualisation, Methodology, Resources, Writing – Review & Editing

**Tom Verguts:** Conceptualisation, Supervision, Funding Acquisition, Writing – Review & Editing

**Gerhard Jocham:** Conceptualisation, Methodology, Supervision, Project administration, Funding Acquisition, Writing – Review & Editing

## Footnotes

1 The model including interactions with the factor Leave (BIC = 262606) did not provide significantly better fit than the reported model (BIC = 262566), xi = 9.93, p = 0.078.

2 For the analysis controlling for the effects of testing location on behaviour, see Supplementary Material 1. These analyses did not reveal significant effects of testing location in behaviour.

