## Supplementary Material for "Patch-leaving decisions and pupil-linked arousal systems"

***1. Controlling for the effects of testing location on behaviour***

In order to ascertain that the behavioural performance did not differ across testing locations, we performed additional analyses including Lab (HHU, UGent) as a fixed effect and tested if including the lab as a predictor improved the model fit.

**Patch residence times.** The following model was performed to evaluate the effect of testing location on patch residence times:

$$Residence \sim Environment+Patch Type+Previous Patch+Environment:Patch Type+Lab +\left( 1 \right| Participant)$$

(S1)

The LME model in Eq. S1 did not significantly differ from the more parsimonious model (Eq. 1), which did not include lab as a fixed effect (*BIC_Eq.S1_* = 11285, *BIC_Eq.1_* = 11277, χ^2^ = 0.065, *p =* 0.798). The main effect of lab was not significant, *β_UGent–HHU_* = –0.22, 95 % CI (-1.93, 1.50), *F*(1,50) = 0.06, *p =* 0.803*,* *η^2^_partial_* = 0.01, 95 % CI (0.00, 1.00), while the significant main effects and interactions reported in the Results all remained significant.

**Overharvesting.** The following model including the effect of testing location was tested:

$$Overharvest \sim Environment+Patch Type+Previous Patch+Environment:Patch Type+Patch Number \left( scaled \right)+Lab+ \left( 1 \right| Participant)$$

(S2 )

This model did not provide a better fit than the more parsimonious model in the Eq. 2 (*BIC_Eq.S2_* = 11728, *BIC_Eq.2_* = 11720, χ^2^ = 0.124, *p* = 0.724*,* and the main effect of lab was not significant, *β_UGent–HHU_* = -0.31, 95 % CI (-2.09, 1.46), *F*(1,50) = 0.12, *p = 0.730,* *η^2^_partial_* = 0.002, 95 % CI (0.00, 1.00). The significant main effects and interactions reported in the Results remained significant.

**RT.** To evaluate the effect of testing location on RT effects, we performed the two following LMEs

$$RT \sim Leave+Environment+Patch Type+Environment:Patch Type+Lab+\left( 1 \right| Participant)$$

(S3)

$$RT on Stay Trials \sim Residence+Environment+Patch Type+Trial Number+Lab+\left( 1 \right| Participant)$$

(S4)

The LME model in Eq. S3 did not significantly differ from the more parsimonious model (Eq. 3), which did not consider lab as a fixed effect (*BIC_Eq.S3_* = 25772, *BIC_Eq.3_* = 25781, χ^2^ = 0.755, *p = 0.385*). The main effect of lab was not significant, *β_UGent–HHU_* = 23.71, 95 % CI (-30.64, 78.07), *F*(1,50) = 0.7312, *p =* 0.397*, η^2^_partial_* = 0.01, 95 % CI (0.00, 1.00), and the main effects and interactions reported in the Results remained significant. Similarly, the model with stay trials as dependent variable, which included lab as a fixed effect (Eq. S4), did not significantly differ from the more parsimonious model (Eq. 4 BIC_Eq. S4_ = 257647, BIC_Eq.4_ = 257638, χ^2^ = 0.815, *p = 0.367*). The main effect of lab was not significant, *β_UGent–HHU_* = 24.59, 95 % CI (–29.65, 78.84), *F*(1,50) = 0.790, *p = 0.378, η^2^_partial_* = 0.02, 95 % CI (0.00, 1.00) and the significant main effects reported in the Results remained significant. In summary, behavioural data did not indicate any significant differences in performance based on the testing location.

***2. Controlling for baseline effects in pupillometry data***

We have analysed the baseline pupil values (averaged from the 500 ms baseline period preceding the stimulus presentation with the following model, identical to in its structure to the model for the analysis of baselined pupil values:

$$Pupil baseline values \sim Leave*Environment*Patch Type+Trial Number \left( scaled \right)+\left( 1 \right|Participant)$$

(S 5)

This analysis has revealed that the baseline pupil values differed as a function of Decision Type, with baseline being smaller for leave compared to stay decisions, *β_Leave–Stay_* = –39.02, 95 % CI (–73.55, –4.47), *F*(1,19755) *=* 63.83, *p*<0.001, *η^2^_partial_ =* 0.003, 95 % CI (0.002 1.00). Baseline pupil values were significantly larger in the HVAR than in the LVAR environment, *β_LVAR–HVAR_* = –64.42, 95 % CI (86.62, 42.22), *F*(1,19754) *=* 10.21, *p* = 0.001, *η^2^_partial_ =* 0.0005, 95 % CI (0.0004, 1.00). The baseline values also varied as a function of patch type, *β_Medium–Small Patch_* = –71.56 , 95 % CI (– 92.52, –50.59), *β_Large–Small Patch_*= –87.48, 95 % CI (– 107.82, –67.15), *F*(2, 19754) *=* 22.92, *p* < 0.001, *η^2^_partial_ =* 0.00009, 95 % CI (0.00, 1.00). The baseline was larger for small reward patches than medium reward patches, estimate = 39.2, SE = 8.54, *t*(19754) = 4.60, *p* < 0.001, and large patches, estimate = 56.5, SE = 8.51, *t*(19754) = 6.64, *p* < 0.001, while the medium and large patch baseline values did not significantly differ, estimate = 17.3, SE = 8.32, *t*(19754) = 2.08, *p* = 0.095. A significant interaction between environment type and patch type, *β_LVARt:MediumPatch_* = 69.61, 95 % CI (41.58, 97.64), *β_LVAR:LargePatch_*= 76.18, 95 % CI (49.01, 103.35), *F*(2, 19754) *=* 14.33, *p* < 0.001, *η^2^_partial_ =* 0.001, 95 % CI (0.0006, 1.00), revealed that the difference between patch baseline values was significant only for the HV environment (small patch in HV vs. medium patch in HV, estimate = 80.32, SE = 12.1, *t*(19754) = 6.65, *p* < 0.001; small patch in HV vs. large patch in HV, estimate = 94.97, SE = 12.0, *t*(19754) = 7.90, *p* < 0.001; while other comparisons were nonsignificant, *p* > 0.212

Baseline pupil values were smaller for HHU participants than UGent participants, *β_UGent – HHU_ =* 1469.71 95 % CI (1129.74, 1809.68), *F*(1,51) = 71.80, *p* < 0.001, *η^2^_partial_ =* 0.58, 95 % CI (0.44, 1.00). Importantly, baseline pupil values significantly decreased as a function of time on task, *β_Trialnumber-scaled_ =* – 23.95, 95 % CI (-28.78, -19.11), *F*(1, 19756) *=* 94.22, *p* < 0.001, *η^2^_partial_ =* 0.005, 95 % CI (0.003, 1.00).

To control for the potential baseline effects in the observed baseline-corrected pupillometry data (cf. Alday, 2019), we also analysed the non-baselined corrected pupillometry data from the time window of the initial dilation (1–2 s), while controlling for the trial-by-trial baseline values with the following LME:

$$Pupil size \sim Leave*Environment Type*Patch Type+Baseline values \left( scaled \right)+Lab+\left( 1 \right| Participant)$$

(S 6)

This model revealed that the observed effect of decision type remained similar (albeit smaller), with leave decisions characterised by a larger pupil dilation than stay decisions, *β_Leave–Stay_* = 43.17, 95 % CI (24.21, 62.13), *F*(1,19760.2) *=* 6.91, *p* = 0.009, *η^2^_partial_ =* 0.0003, 95 % CI (0.00005, 1.00) (see Fig. S1A). The effect of environment was also similar, with pupil dilation being smaller in the LVAR vs HVAR environment, *β_LVAR–HVAR_* = –12.47, 95 % CI (–-24.66, –0.27), *F*(1,19746.2) *=* 25.41, *p* < 0.001, *η^2^_partial_ =* 0.001, 95 % CI (0.0006, 1.00) (see Fig. S1B). The effect of patch type on pupil dilation did not reach significance in this analysis, *β_Medium–Small Patch_* = 3.26 % CI (–8.26, 14.78), *β_Large–Small Patch_*= 5.40, 95 % CI (–5.78, 16.58), *F*(2, 19747.2) *=* 2.43, *p* = 0.088, *η^2^_partial_ =* 0.0002, 95 % CI (0.00, 1.00). See Table S1 for an overview of all effects and interactions.

The results revealed that pupil size in the evidence time window was *smaller* at HHU than at UGent (see Fig S1C), *β_UGent – HHU_ =* 195.74 95 % CI (130.38, 261.10), *F*(1,52.4) = 34.45, *p* < 0.001, *η^2^_partial_ =* 0.40, 95 % CI (0.23, 1.00). The pupil size in the evidence time window was strongly positively correlated with the baseline pupil size values (see Fig. S1D), *β_Baseline-scaled_ =* 815.08, 95 % CI (807.70, 822.45), *F*(1, 17394.3) *=* 46951.51, *p* < 0.001, *η^2^_partial_ =* 0.73, 95 % CI (0.75, 1.00).

| **Interaction effect** | ***β*** | **95 % CI** | **degrees of freedom** | **F** | **p** | **η^2^_partial_** | **95 % CI** |
| --- | --- | --- | --- | --- | --- | --- | --- |
| **Leave:Patch Type** | *β_LeaveDecision::MediumPatch_* = – 29.95  *β_LeaveDecision::LargePatch_* = – 45.84 | (–55.94, –3.96)  (–71.96, –19.99) | 2, 19745.9 | 5.72 | 0.003 | 0.0006 | (0.0001, 1) |
| **Leave: Environment** | *β_LeaveDecision::LVAR_*= – 29.71 | (–56.22,  –3.21) | 2, 19745.9 | 4.21 | 0.041 | 0.0002 | (0.000005, 1) |
| **Environment: Patch Type** | *β_LVARt:MediumPatch_ =* – 2.03  *β_LVAR:LargePatch_ =* 5.47 | (–17.42, 13.37)  (– 9.45, 20.39) | 2, 19744.7 | 3.28 | 0.04 | 00003 | (0.000006, 1) |
| **Leave: Environment:**  **Patch Type** | *β_LeaveDecision:LVARt:MediumPatch_ =* 10.07  *β_LeaveDecision:LVARt:LargePatch_ =* 32.45 | (–26.65, 46.78)  (– 4.12, 69.02) | 2, 19744.7 | 1.56 | 0.202 | 0.0002 | (0.00 1) |

**Table S1 |** Interaction effects for the control analysis of non-baseline corrected pupil size with regressed baseline values.

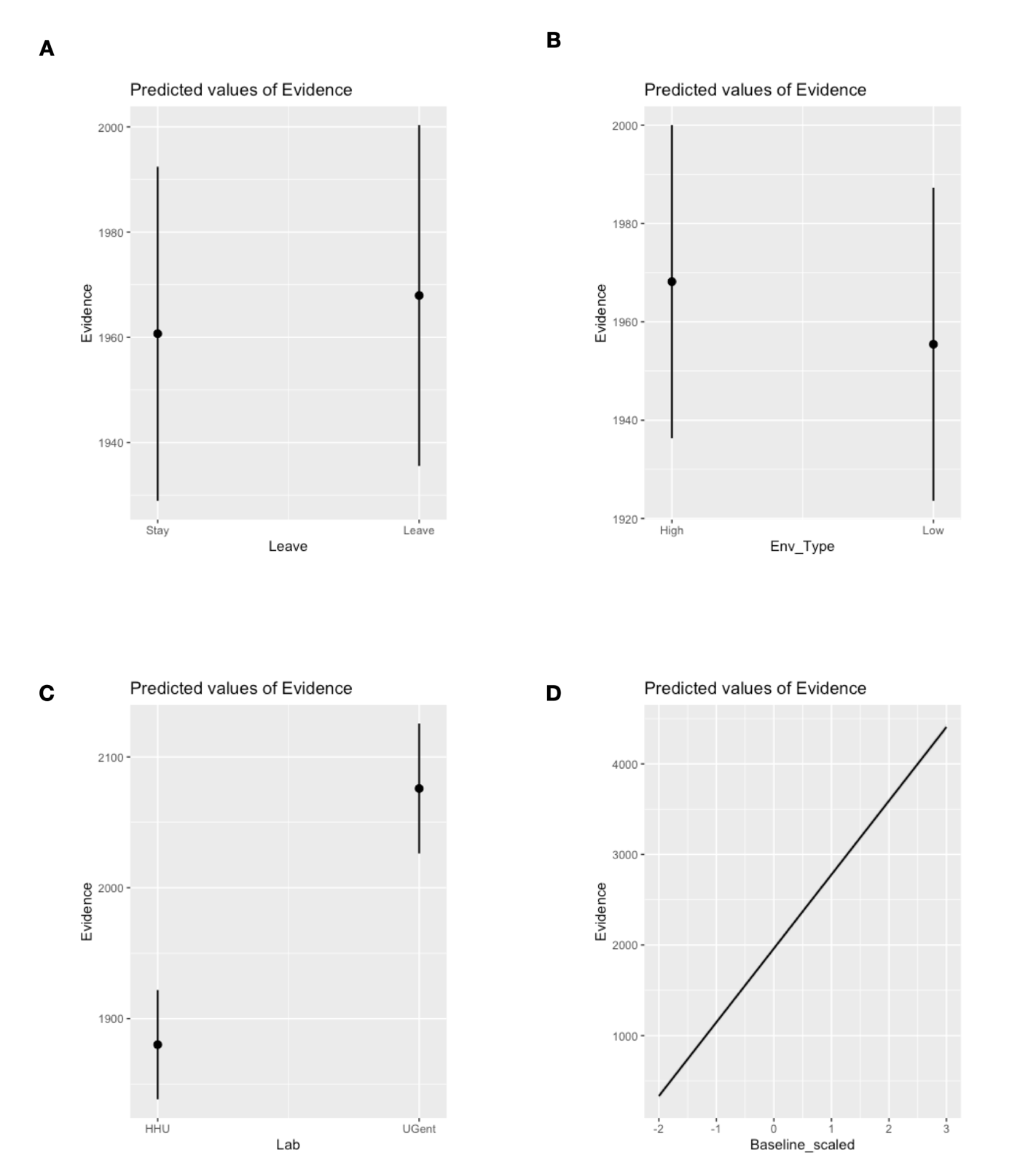
**Figure S1| Regressing baseline values.** Analysis of mean pupil size in the first dilation window (1–2 s) following the evidence with no baseline correction applied (and baseline values regressed): **A.** The effect of decision type: a greater dilation for leave vs. stay decisions. **B.** The effect of environment: larger dilation in the high vs. low variability environment. **C.** The effect of lab: pupil dilation is *smaller* at HHU than at UGent. **D.** Pupil size in the evidence time window is strongly positively correlated with the baseline values.

***3. Time course analysis of pupil derivative***

We additionally performed analyses on the first derivative of the pupil time series, as this measure has been suggested to be less prone to baseline effects (Filipowicz et al., 2020; Fink et al., 2023), and a more accurate predictor of cortical states (Reimer et al., 2016; Yang et al., 2021). We conducted linear mixed effect models on the time series of the whole epoch (-0.499 – 2.5 s) using the following model with pupil derivative values as dependent variables.

$$Pupil derivative \sim Leave+Environment+Environment:Patch Type+Lab+\left( 1 \right|Partcipant)$$

(S7)

The resulting p-values were corrected for multiple comparisons using FDR correction (Benjamini & Hochberg, 1995). We report significant intervals of length of more than 5 ms.

The analysis indicated that the derivative was larger on the trials with leave decisions than on trials with stay decisions (see Fig. 4A). This difference was significant in the following time intervals (FDR-corrected p-values <0.05, interval of adjacent significant values for at least 5 ms): first, in the interval between 326 ms – 684 ms; second, in the interval between 1466 – 2500 ms. Pupil derivatives also differed as a function of patch type (time interval of 5 consecutively significant adjacent time points: 534 – 637 ms) and environment type (1022 –1027 ms). Pupil derivative was also larger for HHU than UGent in the time interval of 542 – 876 ms.

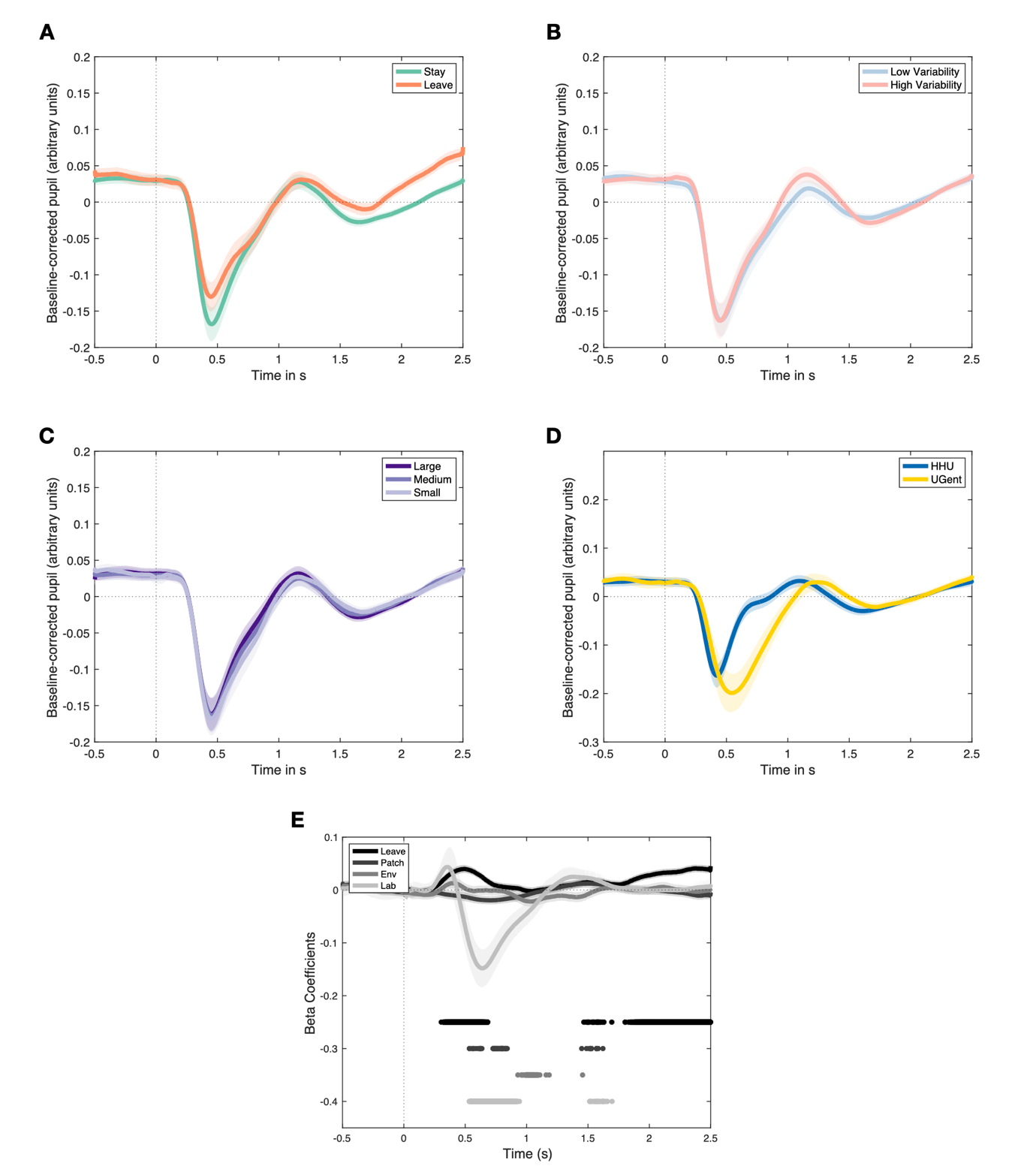

**Figure S2** **| Pupil derivative time series** **A.** Effects of decision type: pupil derivative larger for leave vs. stay decisions. **B**. environmental variability **C.** patch type **D.** testing location **D.** Beta coefficients derived from LME (Eq. S7) on the derivative of pupil time series. Horizontal lines indicate significant FDR-corrected p-values.

***4. Control analysis - value distance from the optimum* reward value at threshold to leave**

We analysed how pupil size varied based on the current reward patch relative to the optimal value of the leave threshold based on the simulations. For this we ran a model with the pupil size from the time window of the initial dilation as a dependent variable, and difference from optimum (quantified as the subtraction of the current patch residence time from the optimal patch residence time), decision Type (leave, stay) as dependent variables, while controlling for the effect of testing location (UGent, HHU) and trial number.

$$Pupil size \sim Difference from Optimum+Leave+Trial Number \left( scaled \right)+Lab+\left( 1 \right|Participant)$$

( S8 )

The analysis showed that pupil size values increased with increasing difference from the optimum, *β_Difference from optimum_ =* 0.12, 95 % CI (0.03,0.21), *F*(1, 19673 ) *=* 7.12, *p* = 0.008, *η^2^_partial_ =* 0.0004, 95 % CI (0.00005, 1.00). The main effect of decision type was significant, *β_Leave–Stay_* = 14.27, 95 % CI (6.11, 22.43), *F*(1,119804) *=* 11.75, *p* < 0.001, *η^2^_partial_ =* 0.0006, 95 % CI (0.0001, 1.00). The main effect of testing location was significant, *β_UGent – HHU_ =* –44.75, 95 % CI (–78.83, –10.67), *F*(1,51) = 6.62, *p* = 0.013, *η^2^_partial_ =* 0.11, 95 % CI (0.01, 1.00). As in other analyses, pupil size was shown to significantly increase with the time on task, *β_TrialNumber-scaled_ =* 5.93, 95 % CI (3.16, 8.71), *F*(1, 19802) *=* 17.56, *p* < 0.001, *η^2^_partial_ =* 0.0009, 95 % CI (0.0003, 1.00).

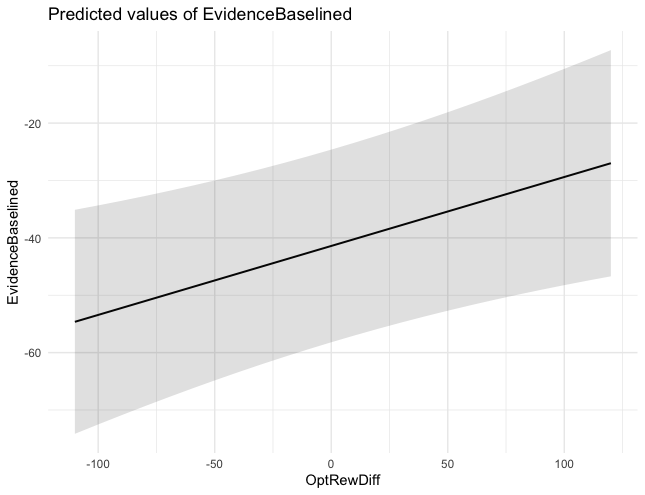

**Figure S3 | Distance from optimum reward threshold at leave.** Analysis of mean pupil size in the first dilation window (1–2 s) following the evidence as a function of distance from optimum reward threshold at leave: Pupil size increases with increasing distance from the optimum.
